# The deacetylase dependent and independent role of HDAC3 in cardiomyopathy

**DOI:** 10.1101/2022.12.23.521758

**Authors:** Jieyu Ren, Qun Zeng, Hongmei Wu, Xuewen Liu, Maria Clara Guida, Wen Huang, Yiyuan Zhai, Junjie Li, Karen Ocorr, Rolf Bodmer, Min Tang

**Affiliations:** Department of Biochemistry and Molecular Biology, College of Hengyang Medical, University of South China, 421001 Hengyang, China; Development Aging and Regeneration Program, Sanford Burnham Prebys Medical Discovery Institute, 10901 N Torrey Pines Rd, La Jolla, CA 92037, USA; Center for Medical Genetics, School of Life Sciences, Central South University, Changsha, 410078 Hunan China

**Keywords:** deacetylase, cardiomyopathy, cardiac contractility, triglycerides, pericardin, Drosophila, fibrosis

## Abstract

**Background:** Cardiomyopathy is a common disease of cardiac muscle that negatively affects cardiac function. HDAC3 commonly functions as co-repressor by removing acetyl moieties from histone tails. However, a deacetylase-independent role of HDAC3 has also been described. Cardiac deletion of HDAC3 causes reduced cardiac contractility accompanied by lipid accumulation. The molecular function of HDAC3 in cardiomyopathy remains unknown. We have used the powerful genetic tools in *Drosophila* to investigate the enzymatic and non-enzymatic roles of HDAC3 in cardiomyopathy.

**Methods and Results:** Using the *Drosophila* heart model, we showed that cardiac-specific HDAC3 knockdown leads to prolonged systoles and reduced cardiac contractility. Immunohistochemistry revealed structural abnormalities characterized by myofiber disruption in HDAC3 knock down hearts. Cardiac-specific HDAC3 knockdown showed increased levels of whole body triglycerides and increased fibrosis. The introduction of deacetylase-dead HDAC3 mutant in HDAC3 KD background showed comparable results with wild type HDAC3 in aspects of contractility and Pericardin deposition. However, deacetylase-dead HDAC3 mutants failed to improve triglyceride accumulation.

**Conclusions:** Our data indicate that HDAC3 plays a deacetylase-independent role in maintaining cardiac contractility and preventing Pericardin deposition as well as a deacetylase-dependent role to maintain triglyceride homestasis.

## Introduction

Cardiomyopathies are complex disorders that arise from a heterogeneous group of pathologies in cardiomyocytes. At the cellular level, cardiomyopathies are characterized by structural and functional changes that include imbalances in metabolic pathways [1], and extracellular matrix (ECM) deposition [2]. Histone deacetylases (HDACs) are enzymes that modulate gene transcription and have been shown to play roles in cardiac hypertrophy [3]. For example, mice with HDAC2 overexpression are sensitive to hypertrophic stimuli, whereas mice lacking HDAC2 are resistant to hypertrophic stress [4]. On the other hand, mice lacking HDAC5 and HDAC9 are sensitized to cardiac stress signals and develop severe cardiac hypertrophy in response to pressure overload by inducing the re-expression of fetal genes [5], suggesting the different roles for HDACs in cardiomyopathy.

HDAC3 have been shown to play unique roles in maintaining cardiac metabolic balance. MCK-Cre mediated cardiac deletion of HDAC3 after birth has no obvious effect on cardiac function. However, severe cardiac hypertrophy was found in postnatal HDAC3 depleted mice upon switching to a high fat diet, and was associated with decreased the expression of fatty acid oxidation enzymes [6]. Earlier cardiac-specific deletion of HDAC3 during embryo development by MHC-αCre resulted in cardiac hypertrophy and fibrosis, accompanied by up-regulation of genes involved in fatty acid uptake and oxidation [7].

HDAC3 belongs to class I HDACs and exists as part of a complex that contains either nuclear receptor corepressor (NCoR) or its homolog silencing mediator of retinoic and thyroid receptors (SMRT) [8, 9] and it is activated by physical interactions with the conserved domain in NCoR and SMRT, deacetylase activating domain (DAD) [9, 10]. HDAC3 commonly functions as a co-repressor by removing acetyl moieties from histone tails. However, a deacetylase-independent role of HDAC3 has also been identified. Global deletion of HDAC3 is embryonic lethal, whereas knock-in mice bearing the mutations in the DADs of both NCoR and SMRT (NS-DADm) live to adulthood despite undetectable deacetylase activity of HDAC3 in the embryo [11, 12]. Furthermore, the non-enzymatic activity of HDAC3 silences the cardiac lineage genes through targeting to the nuclear lamina to repress cardiac progenitor differentiation [13]. HDAC3 also coordinates deacetylase-independent epigenetic silencing of Transforming Growth Factor-β1 to prevent the development of cardiac fibrosis [14]. Deacetylase-dead mutant can partially rescue HDAC3-dependent phenotypes in the mouse liver [15]. HDAC3 is necessary for epidermal stratification independent of its deacetylase activity [16] However, the catalytic function of HDAC3 in cardiac performance remains unknown. Here we take advantage of *Drosophila* cardiac model and used deacetylase-dead transgenic mutant to investigate the enzymatic and non-enzymatic roles of HDAC3 in cardiomyopathy.

## Result

### 1. HDAC3 is required to maintain adult heart physiological function

Cardiac specific HDAC3 knock down is achieved by crossing heart specific driver hand4.2 Gal4 with RNA interference line with short hairpin RNA targeting HDAC3 (Short hairpin ID: SH00372.N), referred as HDAC3 KD. Quantitative polymerase chain reaction showed that HDAC3 mRNA expression in the HDAC3 KD hearts was 52.3% of the level expressed in control hearts (Supplemental figure 1).

**Figure 1.**
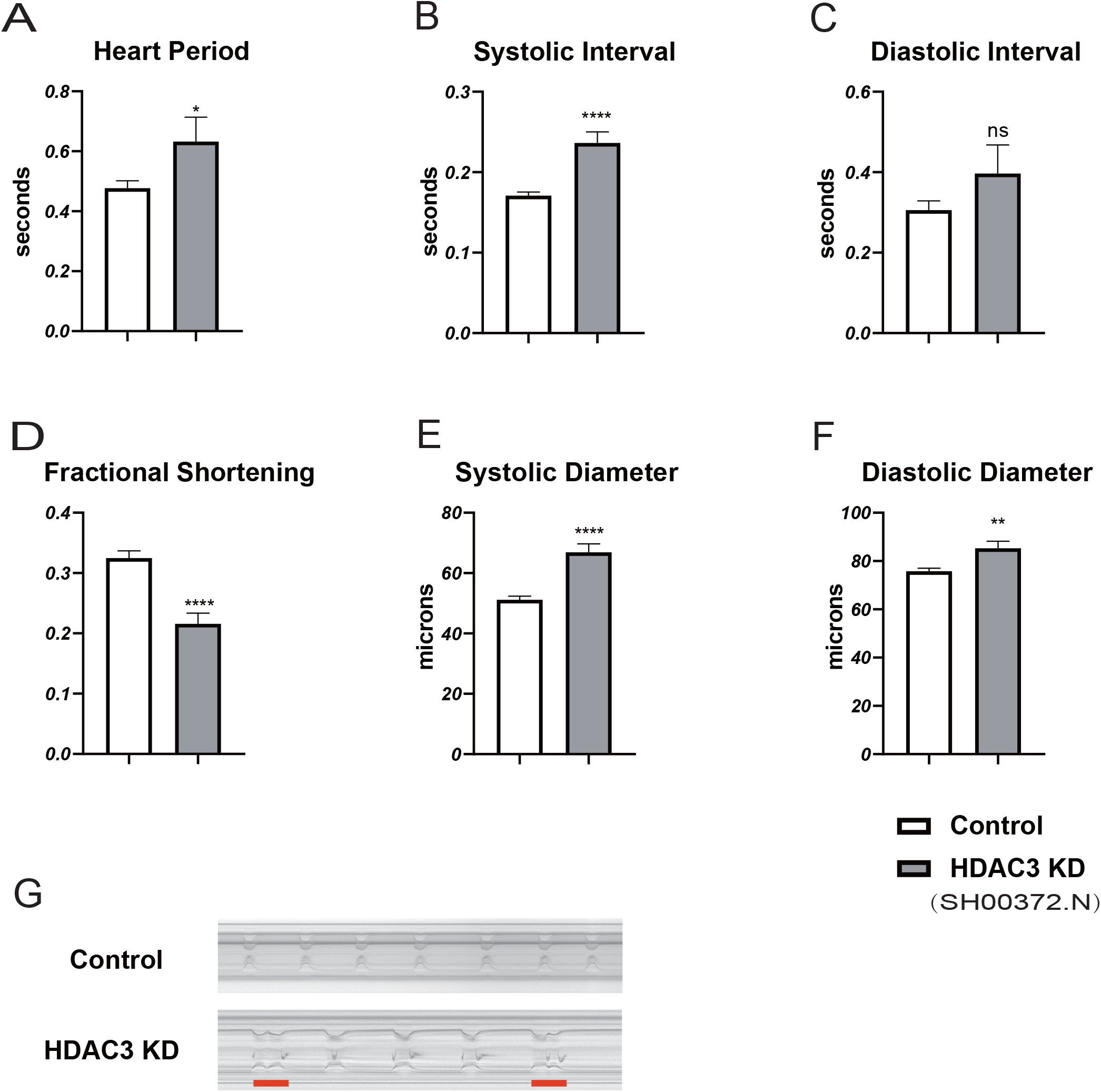
Heart specific HDAC3 KD causes cardiac physiological dysfunction. (A) Heart Period, (B) Systolic Interval, (C) Diastolic Interval, (D) contractility quantified as Fractional Shortening, (E) Systolic Diameter, and (F) Diastolic Diameter were measured for hearts from wild type control flies (the progeny of the crosses of Hand-Gal4 x UAS-control line, and flies with cardiac specific HDAC3 knockdown (Hand>UAS-HDAC3 RNAi, SH00372.N). (G) Five-second M-modes from movies of control, HDAC3 KD. Fibrillation in HDAC3 KD is highlighted with red line. Note the significant systolic interval prolongation and decreased contractility in HDAC3 KD. Significance was determined using unpaired student’s t tests. Differences relative to the control are indicated by individual asterisks; Graph Pad statistical analysis, *P<0.05, **P<0.01, ****P<0.0001. ns represents no significance. Sample size was 20 to 30 flies per genotype.

The cardiac phenotype induced by HDAC3 KD is illustrated in high-speed movies using a transgenic heart marker R94C02::tdTomato[17], which show irregular heart beats and structural abnormalities (Supplemental Movie 1 & 2). We characterized the effects of cardiac-specific HDAC3 knockdown on cardiac function using the Semi-automated optical heart beat analysis system [18]. Cardiac-specific HDAC3 KD leads to prolonged heart periods (Figure 1A), which is due to prolonged systolic intervals (Figure 1B). Knockdown of HDAC3 also affected cardiac contractility, as demonstrated by decreased Fractional Shortening (Figure 1D). The cardiac phenotype induced by HDAC3 KD is also illustrated in the M-mode traces obtained from high-speed movies, which show heart wall movements over time. M-Modes from HDAC3 KD hearts exhibited long pauses between beats (asystoles), prolonged or multiple (fibrillatory) contractions (Figure 1G).

The percentage of prolonged systoles (>0.4 seconds, the average systolic interval of wild-type flies range from 0.15 to 0.25), reminiscent of unsustained fibrillations, was markedly elevated in HDAC3 KD flies compared with controls (Table 1). The percentage of prolonged diastoles (>1 seconds, the average systolic interval of wild-type flies range from 0.2 to 0.6), indicative of asystolic events, also increased in HDAC3 KD hearts (Table 1). Consistently, the percentage of flies with prolonged diastoles or systoles also increased in HDAC3 KD.

**Table 1.** The prevalence of prolonged systoles and Diastoles in HDAC3 KD.

|  | **% Long**<br>**Diastoles**<br>**(>1s)** | **% Long**<br>**Systoles**<br>**(>0.4s)** | **%Flies**<br>**with DI > 1s** | **%Flies**<br>**with SI >**<br>**0.4s** |
| --- | --- | --- | --- | --- |
| **control** | **0.00%** | **0.00%** | **5.88%** | **5.88%** |
| **HDAC3 KD**<br>**(SH00372.N)** | **1.82%** | **3.90%** | **19.23%** | **34.62%** |
| **HDAC3 KD**<br>**(SH003161.N)** | **0.50%** | **2.76%** | **16.67%** | **20.83%** |

To confirm the phenotype observed in HDAC3 KD which was not due to off-target effects, another independent HDAC3 KD line (Short hairpin ID: SH003161.N) were observed and displayed a similar phenotype demonstrated by prolonged systolic interval and reduced contractility (Supplemental figure 2 and Table 1).

**Figure 2.**
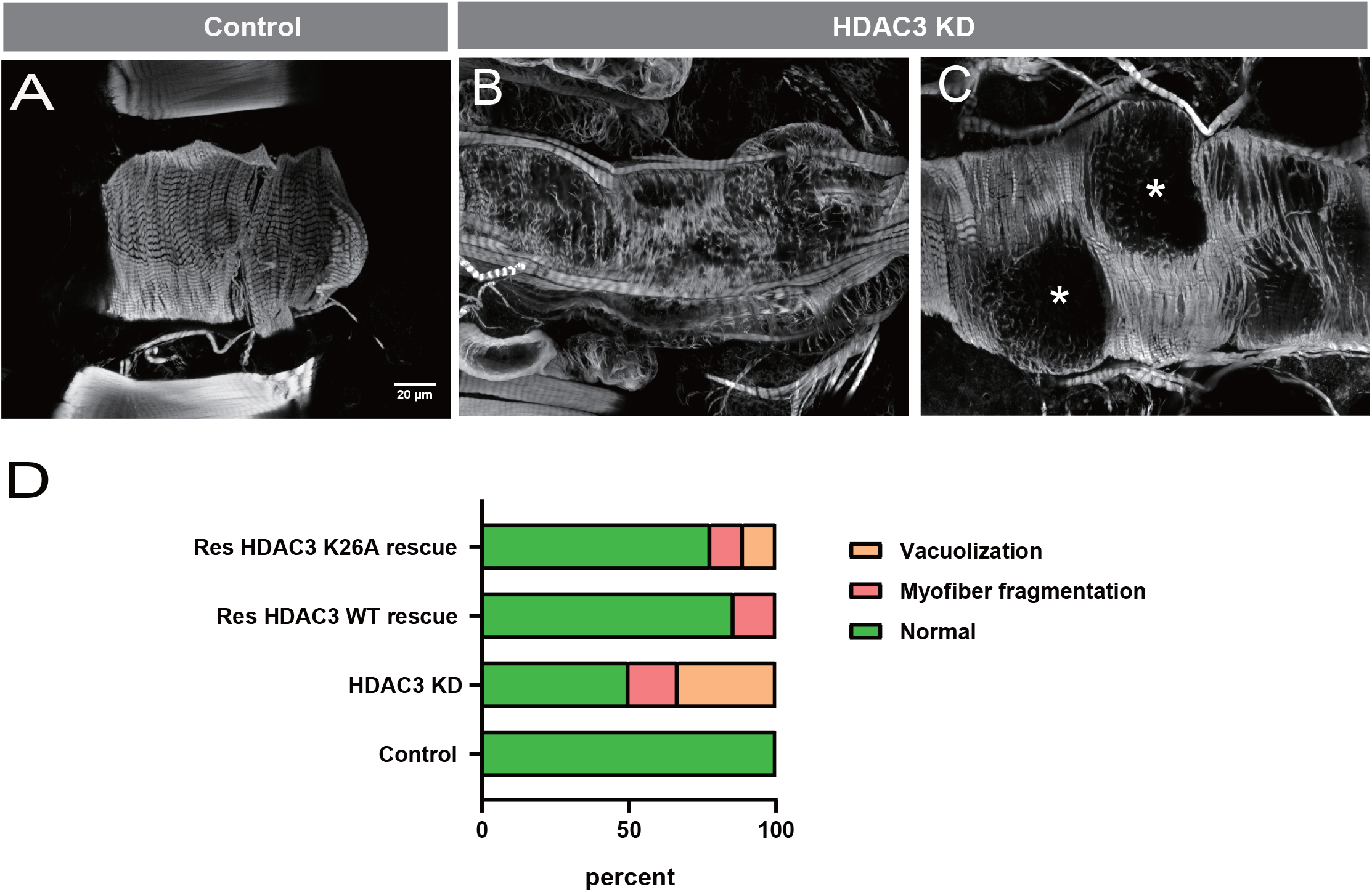
Heart specific HDAC3 knockdown compromises structural integrity. (A-C) Phalloidin staining for sarcomeric actin filaments in cardiomyocytes. Anterior is left in all images. (A) Cardiomyocytes from wild-type controls (Hand-Gal4 > UAS-control line) contain densely packed and circumferentially organized myofibrils. Knock down of HDAC3 fragmented myofibrils (B) and even caused vacuoles formation within cardiomyocytes (asterisk) (C). (D) The percentage of cardiomyocytes with normal (green), myofibril fragmentation (pink), or vacuolization (orange) in control, HDAC3 KD, wild type (wt) HDAC3 rescue flies (hand>UAS-HDAC3 RNAi; shRNA-resistant wt HDAC3 transgene), and HDAC3 K26A mutant rescue flies (hand>UAS-HDAC3 RNAi; shRNA-resistant HDAC3 K26A transgene). More than 20 hearts was observed per genotype.

### 2. HDAC3 KD disrupted myofibrils intergrity

The reduction in cardiac contractility with HDAC3 KD suggested that cardiac morphology in HDAC3 KD flies might also be changed. Control hearts stained for F-actin with Phalloidin showed densely organized myocardial fibers organized circumferentially around the heart tube. In contrast, HDAC3 KD hearts exhibited morphological abnormalities, demonstrated by fragmented myofibrils (Figure. 2B) and even the vacuoles forming (Figure. 2C) within cardiomyocytes.

### 3. Cardiac specific HDAC3 KD increased fibrosis

An accumulation of extracellular matrix protein, such as type-IV-like collagen (Pericardin), has been linked to reductions in cardiac contractility in *Drosophila*, and is reminiscent of cardiac fibrosis [19]. Immunofluorescence staining with Pericardin antibodies revealed an increase of the extracardiac collagen at the distal heart region in HDAC3 KD (Figure 3A-B). The measurement of immunofluorescence intensity also showed increased of Pericardin in response to cardiac HDAC3 KD (Figure 3E). Accordingly, transcript levels of *pericardin* was significantly increased (Figure 3H). Western blotting of lysates of isolated hearts further confirmed an increased level of Pericardin in HDAC3 KD hearts (Figure 3F-G).

**Figure 3.**
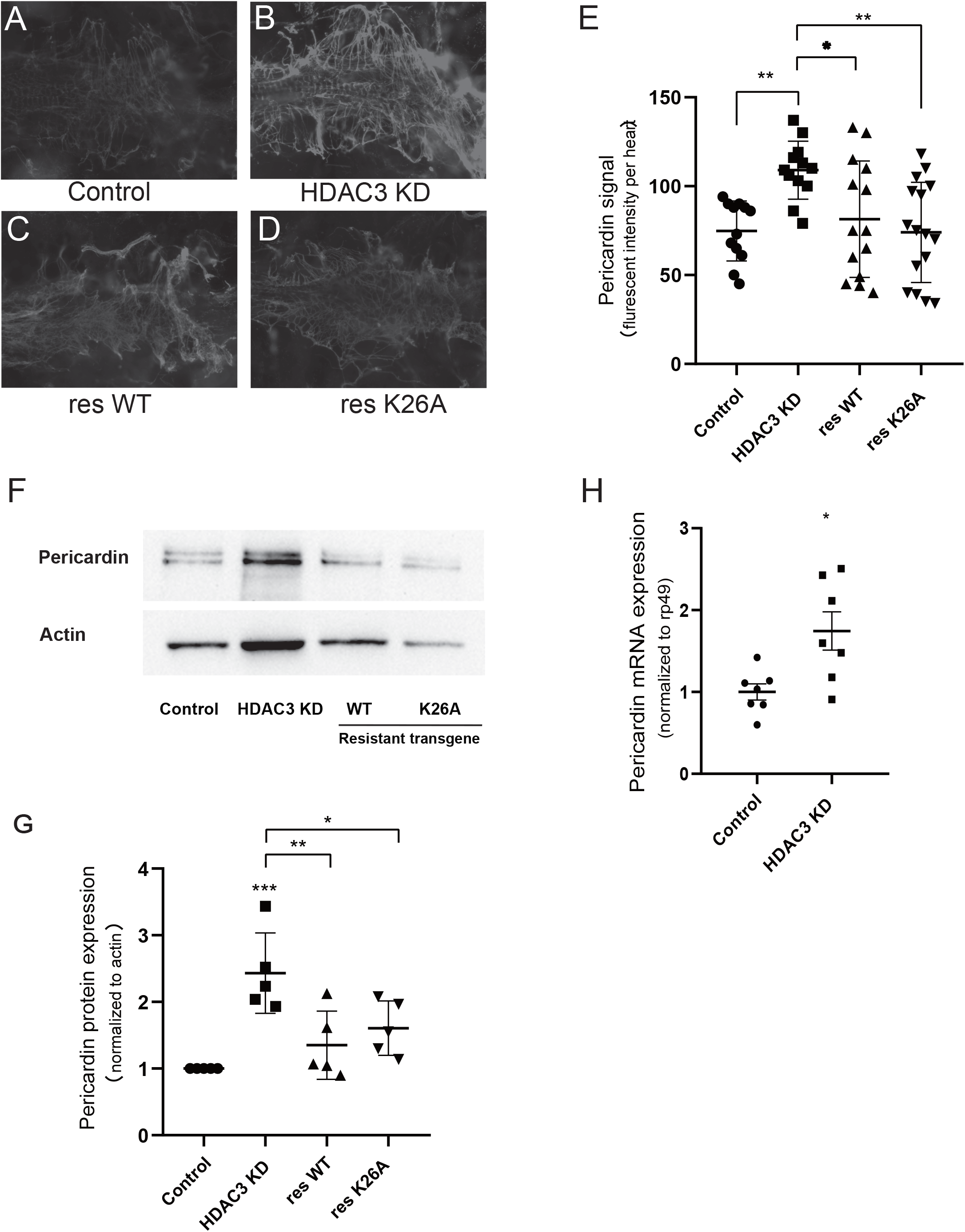
Deacetylase-dead HDAC3 mutant rescued the increased Pericardin deposition in heart specific HDAC3 KD. Pericardin staining at the distal heart of Drosophila from a control (A), HDAC3 KD (B), HDAC3 WT rescue flies (C), and HDAC3 K26A mutant rescue flies (D). Anterior is to the left in all images. (E) Quantification of Pericardin fluorescent signal at the distal heart. (G) Western blot of heart lysates stained with anti-pericardin and anti-actin (used as a loading control). (F) Quantification of Pericardin band intensity relative to actin using Image J software. n=5. (H) Relative mRNA expression of *pericardin* in hearts was normalized to ribosomal rp49 expression. n=7. Note that HDAC3 KD showed increased Pericardin expression, which could be rescued by shRNA-resistant wt or deacetylase-dead mutant HDAC3 transgene. Significance for Pericardin fluorescent signal and Pericardin protein expression was determined using 1-way ANOVA. Significance for *pericardin* mRNA expression was determined using unpaired T-test. Differences among each genotype are indicated by individual asterisks; Graph Pad statistical analysis, *P<0.05, **P<0.01, ***P<0.001. ns represents no significance.

### 4. HDAC3 KD increased levels of triglycerides in flies

Excessive accumulation of lipids is a high risk factor for cardiomyopathy [20, 21]. HDAC3 plays a distinct role in maintaining myocardical lipid homeostasis compared to class I HDACs, HDAC1 and HDAC2 in mice [7, 22]. We quantified whole body triglyceride levels in female flies with cardiac HDAC3 KD. We observed a dramatic increase in myocardial triglycerides (TAG) in HDAC3 KD (Figure. 4A). The increase of triglycerides in HDAC3 KD hearts is comparable with hearts from wild type flies fed with high-fat diet containing 30% coconut (Figure. 4A). Interestingly, cardiac HDAC3 KD also induced an increase in whole body triglyceride content (Figure. 4B).

**Figure 4.**
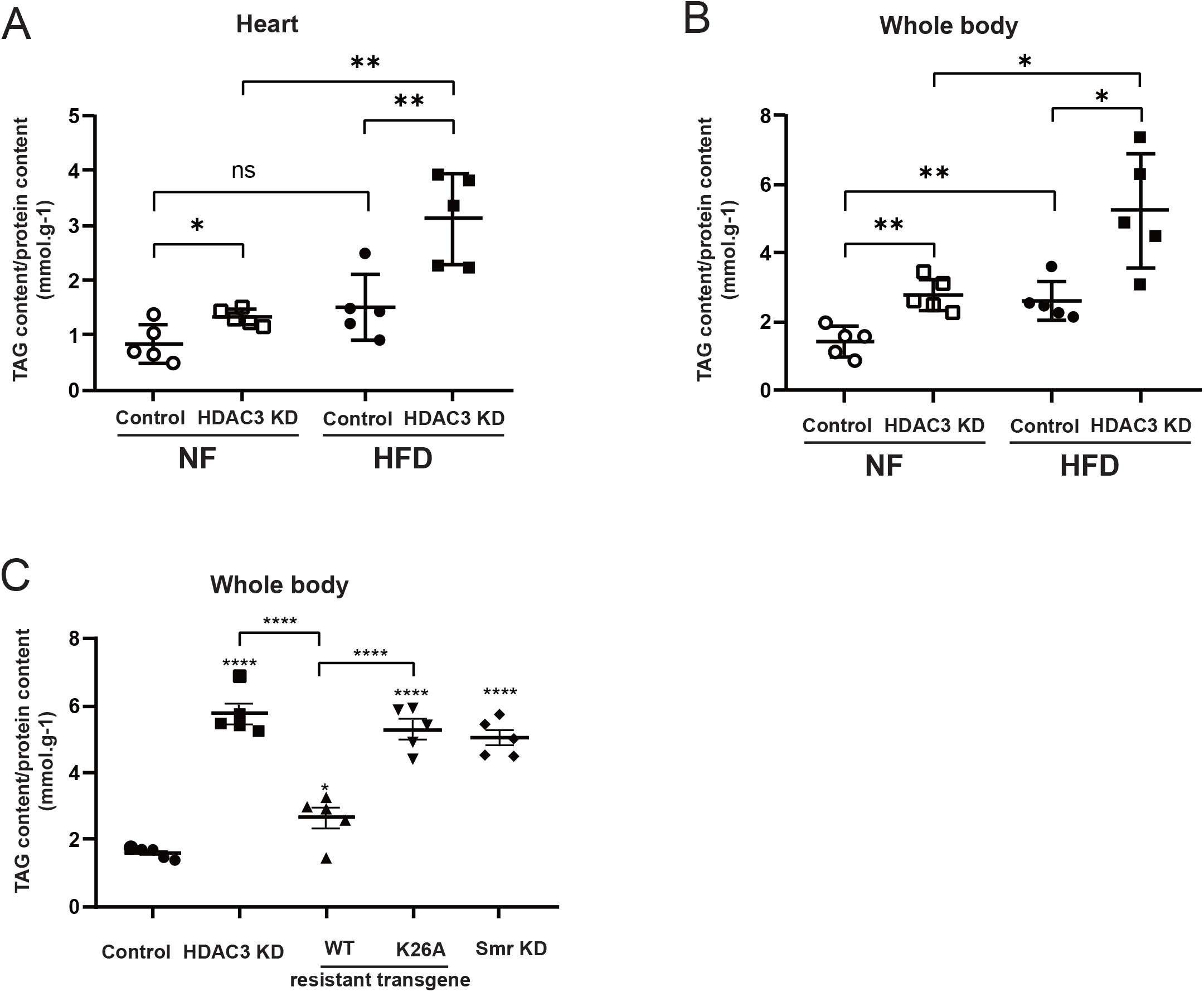
The deacetylase dependent role of HDAC3 is required for regulating TAG levels. (A) Relative TAG content (normalized to protein content) of female hearts on standard diet and high fat diet.. (B) Relative TAG content of the whole body of female flies in control and HDAC3 KD under NF or HFD condition. (C) Relative TAG content (normalized to protein content) of the whole body of female flies in control, HDAC3 KD, HDAC3 WT rescue flies, HDAC3 K26A mutant rescue flies, and Smr KD under NF condition. Note that Cardiac HDAC3 KD caused dramatic increases in TAG levels compared to controls under both NF and HFD conditions. High-fat diet has additive effect on triglycerides accumulation of HDAC3 KD. The accumulation of TAG in whole body was rescued by shRNA-resistant HDAC3 WT transgene but not by HDAC3 K26A mutant. Smr KD showed significant increases in TAG levels comparable to that for HDAC3 KD flies. Significance was determined using 1-way ANOVA (Figure C) or 2-way ANOVA (Figure A-B). Significance between experimental groups is indicated by the capped lines; Differences relative to the control are indicated by asterisks; Graph Pad statistical analysis, *P<0.05, **P<0.01, *** P<0.005, **** P<0.001. ns represents no significance. 5 biological sample per genotype.

HDAC3 can form a complex with NCoR and SMRT in mammals, and its enzymatic activity requires interaction with NCoR/SMRT [8, 10]. Drosophila Smr is homologous to mammalian NCoR and SMRT genes. Cardiac Smr KD showed similar increases in TAG content, consistent with the effects of HDAC3 KD (Figure. 4C).

### 5. Deacetylase dead HDAC3 mutants rescue both cardiac physiological and morphological defects

The crystal structure of HDAC3 revealed that K25 residue is a key residue that contacts the DAD domain of NCoR/SMRT [9, 10, 23]. HDAC3 harboring a K25A mutation disrupted the deacetylase activity[12].The corresponding HDAC3 K26A mutation in *Drosophila* also disrupted deacetylase activity [24]. To investigate if the cardiac phenotype in HDAC3 KD is dependent on the deacetylase activity, we generated wild type or deacetylase-dead mutant HDAC3 transgenic flies under its own endogenous promoter. Deacetylase-dead mutation K26A was introduced in HDAC3 deacetylase-dead mutant. Additionally, the trangenic gene was mutated at RNA interference targeting sites by using synonymous coden to resist shRNA targeting, so referred as shRNA-resistant transgene [24].

The introduction of one copy of deacetylase-dead HDAC3 transgene in HDAC3 KD background showed comparable results with the introduction of wild type HDAC3 transgene in aspects of systolic interval and contractility (Figure 5A-B). Phalloidin staining also showed that deacetylase-dead HDAC3 K26A mutant rescued cardiac morphology abnormalities in the HDAC3 KD, which is comparable with wild type HDAC3 (Figure 2D). In addition, deacetylase-dead HDAC3 K26A mutant significantly reduced the fibrosis associated with HDAC3 KD (Figure.3D-F). Taken together, deacetylase-dead HDAC3 K26A was able to rescue the functional and morphological defects in HDAC KD hearts.

**Figure 5.**
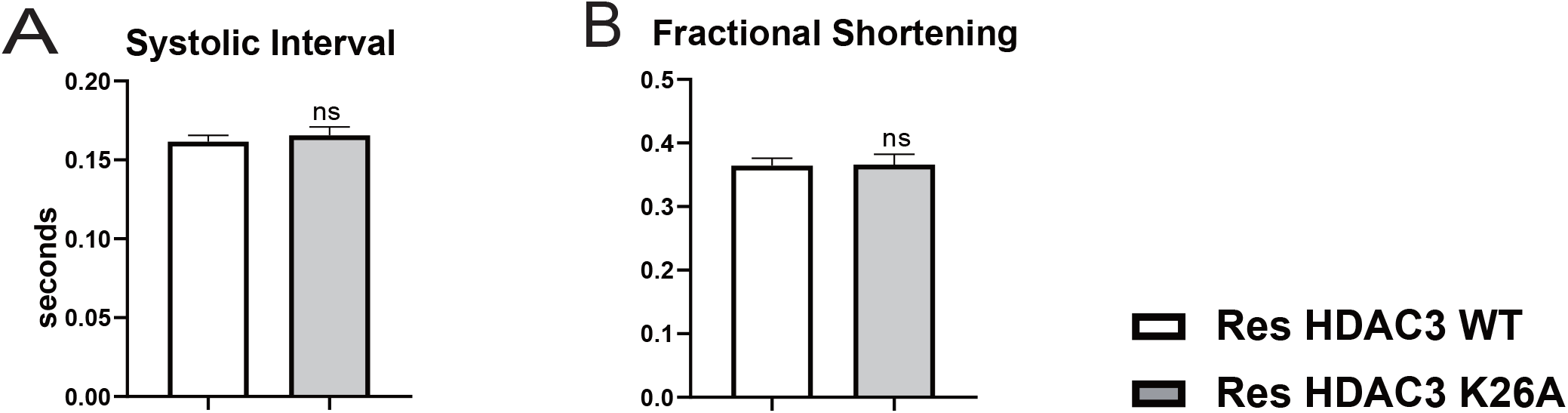
shRNA-resistant deacetylase-dead HDAC3 mutant in HDAC3 KD showed comparable result with wild type HDAC3. (A) Systolic Interval and (B) contractility quantified as Fractional Shortening, were measured for HDAC3 WT rescue flie and HDAC3 K26A mutant rescue flies. Note deacetylase-dead HDAC3 K26A rescue flies showed comparable results with wild type HDAC3 in aspects of heart period, systolic interval, and contractility. Significance was determined using unpaired student’s t tests. Differences relative to the control are indicated by individual asterisks; Graph Pad statistical analysis, *P<0.05, **P<0.01, ns represents no significance. Sample size was 20 to 30 flies per genotype.

### 6. Deacetylase dead HDAC3 mutants failed to improve triglycerides accumulation in HDAC3 KD

HDAC KD caused TAG accumulation of whole body that could be almost rescued by shRNA resistant wild type HDAC3 (Fig. 4C). However, deacetylase-dead HDAC3 K26A mutant failed to rescue the elevated TAG content, indicating the regulation of TAG level by HDAC3 is deacetylase dependent (Figure. 4C).

### 7. HDAC3 KD decreased the median survival time

The heart function is a key factor for the longevity. To test the effects of cardiac HDAC3 KD on the longevity, we aged cardiac specific HDAC3 knockdown flies and wild type control flies in groups of 20 flies each on standard food at 21°C. Cardiac specific HDAC3 KD decreased the median survival time of the flies by 42% (Figure.6).

### 8. High fat diet feeding did exacerbate the effects of HDCA3 KD on longevity reduction but not on cardiac contractility and Pericardin deposition

High fat diet (HFD) is a high risk factor in causing longevity and cardiac dysfunction [25], which led us to test if high fat diet have additive effect on HDAC3 KD phenotype. High fat-diet resulted in a further increase of TAG content in HDAC KD (Figure. 4A-B). Survival assay showed that high-fat diet further decreased the median of survival time in HDAC3 KD (Figure. 6). We also examined the heart function in wild type control flies and HDAC3 KD flies fed HFD. As expected, hearts from control flies fed HFD exhibited a reduction in fractional shortening. Contractility in the HDAC3 KD hearts was not worsened upon HFD feeding (Figure. 7A). Consistently, HFD feeding would not further increase the Pericardin deposition (Figure. 7B).

**Figure 6.**
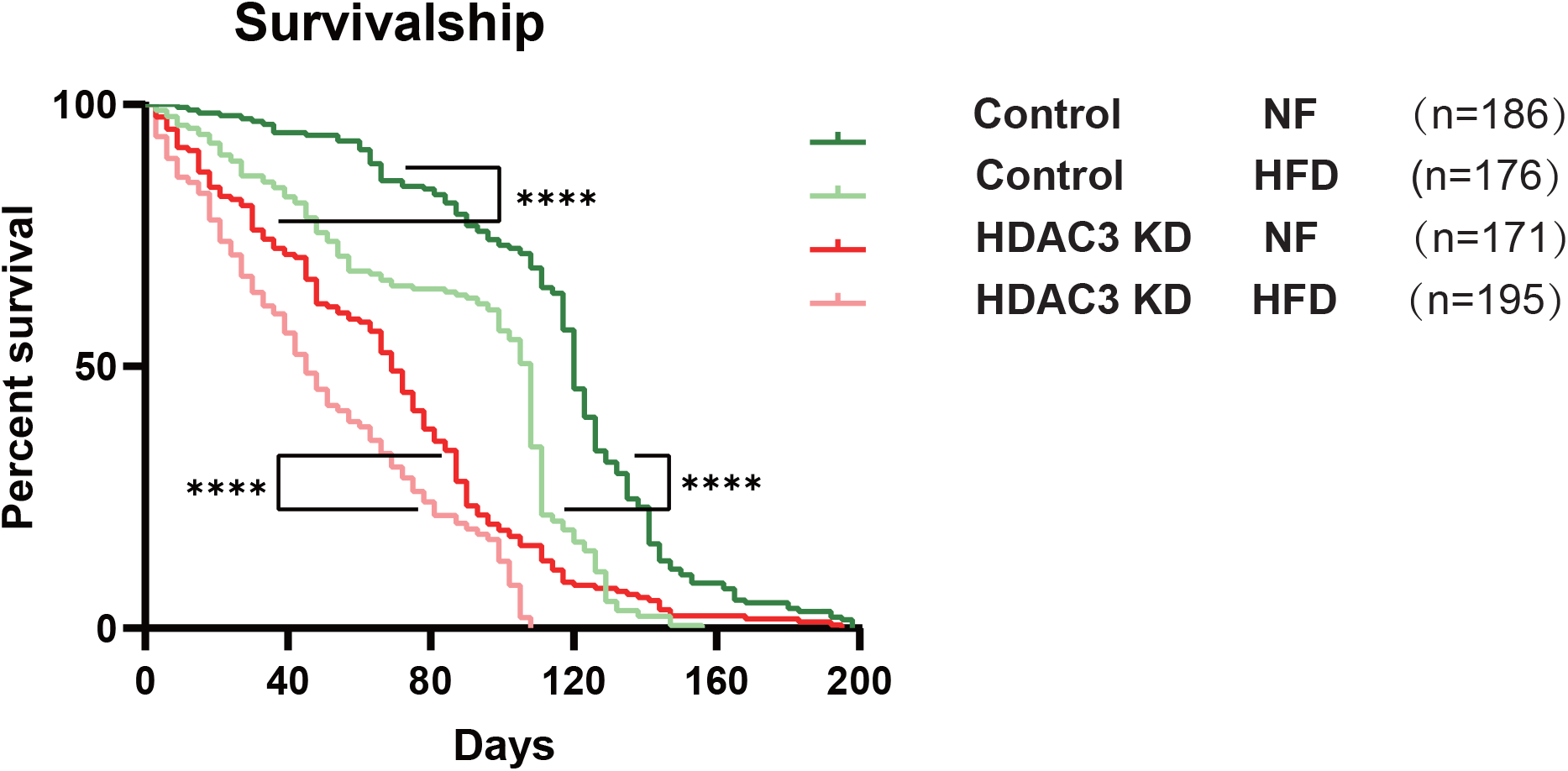
Cardiac specific HDAC3 KD reduces life span. Median survivorship was 120 days for control flies (Hand > UAS-control) compared to 69 days for cardiac specific HDAC3 KD flies (Hand > UAS-HDAC3 RNAi) under NF conditions. Under HFD conditions the median survival of was significantly reduced in controls (to 108 days, a 10% reduction) as well as in HDAC3 KD flies (to 45 days, a 35% reduction). Graph plots % survival versus time (in days) post-eclosion (****p<0.0001, Mandel-Cox log rank test).

**Figure 7.**
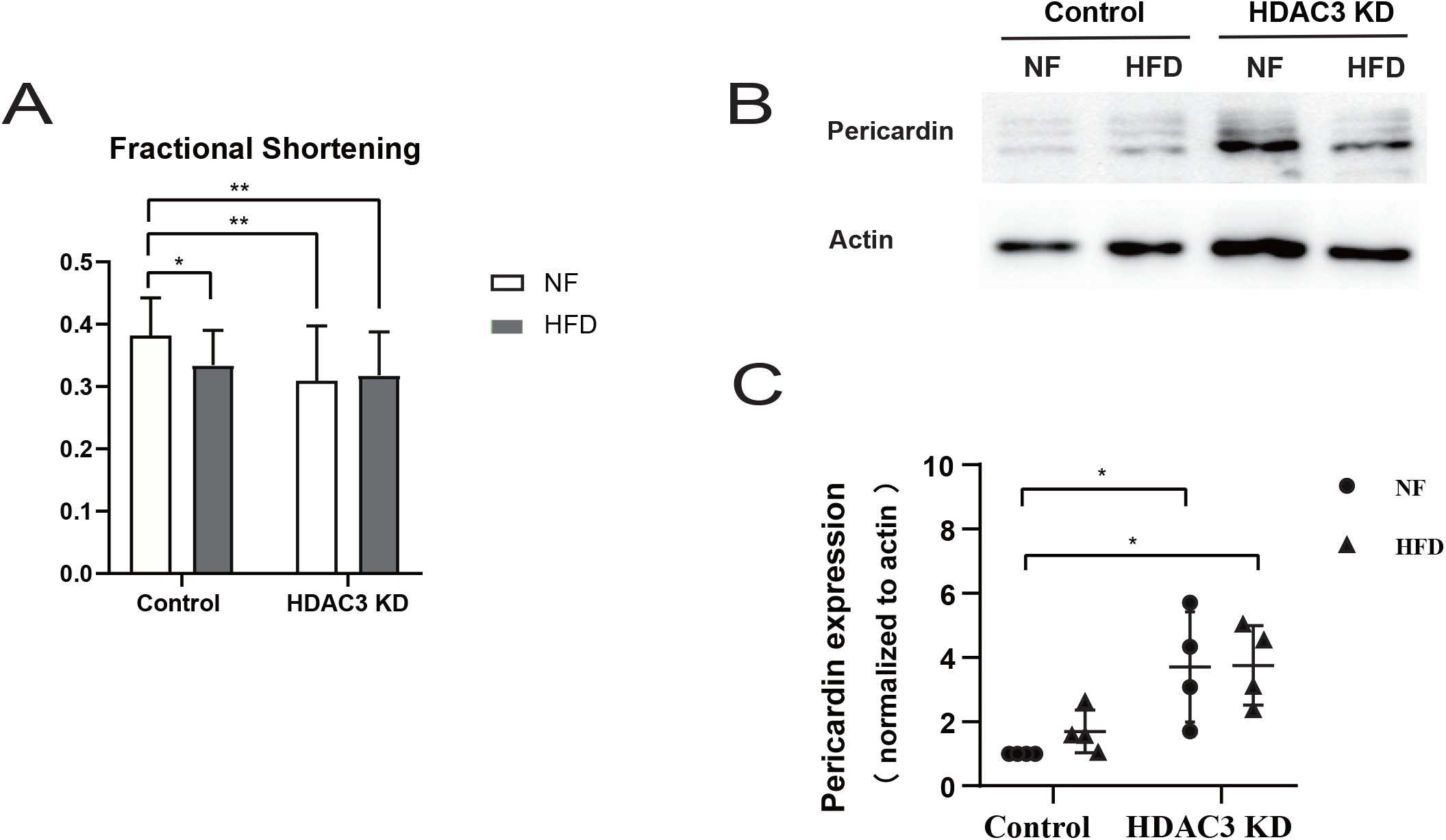
HFD feeding did not exacerbate the effects of HDCA3 KD on cardiac contractility and Pericardin deposition. (A) Cardiac contractility quantified as fraction shortening in control and HDAC3 KD during NF and HFD condition. Sample size was 20 to 30 flies per genotype. (B) Western blot of heart lysates stained with anti-pericardin and anti-actin (used as a loading control). (G) Quantification of Pericardin band intensity relative to actin using Image J software. Note that cardiac HDAC3 KD caused a significant reduction in cardiac contractility and increase in Pericardin deposition. HFD failed to exist additional effect in HDAC3 KD hearts. Significance was determined using 2-way ANOVA. GraphPad statistical analysis, *p<0.05, **p<0.01.

## Discussion

The results described here provide strong evidence that HDAC3 is required to maintain cardiac physiological function and structural integrity in *Drosophila*, demonstrated by reduced cardiac contractility, disrupted myofibrils, and fibrosis in response to cardiac-specific HDAC3 KD, which is reminiscent of aspects of cardiomyopathies in humans and in mice [7], indicating the conservation of HDAC3 function in maintaining adult heart physiology and structure.

The HDAC3 KD induced reduction in cardiac contractility was consistent with the observed disruption of cardiac structure. In addition, the accumulation of ECM protein Pericardin was observed in HDAC3 KD hearts. ECM protein surround the lateral surfaces of cardiomyocytes and accumulate in the ageing heart, which accompanied reduced contractility, referred as cardiac fibrosis [2, 19]. The excessive deposition of ECM proteins in HDAC3 KD likely contributed to the decreased cardiac contractility. HDAC3 has been shown to recruit the PRC2 complex to epigenetic silence transforming growth factor-β signaling[14]. TGF-βsignaling play a central role in cardiac fibrosis [26, 27]. So we speculated that HDAC3 is required to prevent cardiac fibrosis through inhibiting TGF-βsignaling.

HDAC3 play a distinct role in cardiac metabolism compared to other class I HDACs [7, 22]. HDAC3 participates in fatty acid (FA) oxidation [6, 28] and the defect of FA oxidation would be expected to lead to lipotoxicity, accompanied with the increased TAG levels. As expected, cardiac specific HDAC3 KD increased the TAG levels. Similarly, loss of HDAC3 caused hepatic steatosis by rerouting metabolites towards lipid synthesis and inhibiting fatty acid oxidation [29]. Bradycardia in HDAC3 KD would be expected due to reduced supply of the energy substrate that reroutes to TAG synthesis.

Cardiac HDAC3 KD induced an increase of triglyceride content in whole body. The heart, as a major energy consuming organ, would be expected to have an impact on the whole body TAG in *Drosophila* with low metabolite rate. Additionally, it also would be possible that HDAC3 may regulate the non-autonomous crosstalk factors, such as metabolites, which may contribute to the changes of TAG levels in the whole body. The deacetylase-dead mutant K26A could rescue the contractility and cardiac fibrosis -like phenotypes, indicating the role of HDAC3 in these aspects of cardiac function is deacetylase independent. Consistent with our observations, previous studies have shown that HDAC3 plays a deacetylase-independent role in inhibiting extracellular matrix scaffold protein expression during mouse heart development, mediated by the transforming growth factor-β pathway [14]. Consistently, deacetylase-dead mutation K26A still maintain the repressor activity of HDAC3 during embryo development in *Drosophila* [24]. So we speculated that the non-enzymatic role of HDAC3 is involved in maintaining adult heart function through inhibiting ECM protein Pericardin in *Drosophila*.

The deacetylase-dead mutant K26A failed to rescue the increased whole body TAG levels in HDAC3 KD, indicating that the regulation of TAG levels is dependent on acetylation activity. This result was confirmed by KD of Smr, the homologue of mammalian NCoR/SMRT that is required for the enzymatic activity of HDAC3 [9, 10]. Together, our result showed the deacetylase-dependent role of HDAC3 in lipid homeostasis and deacetylase-independent role in ECM accummulation (Figure. 8).

**Figure 8.**
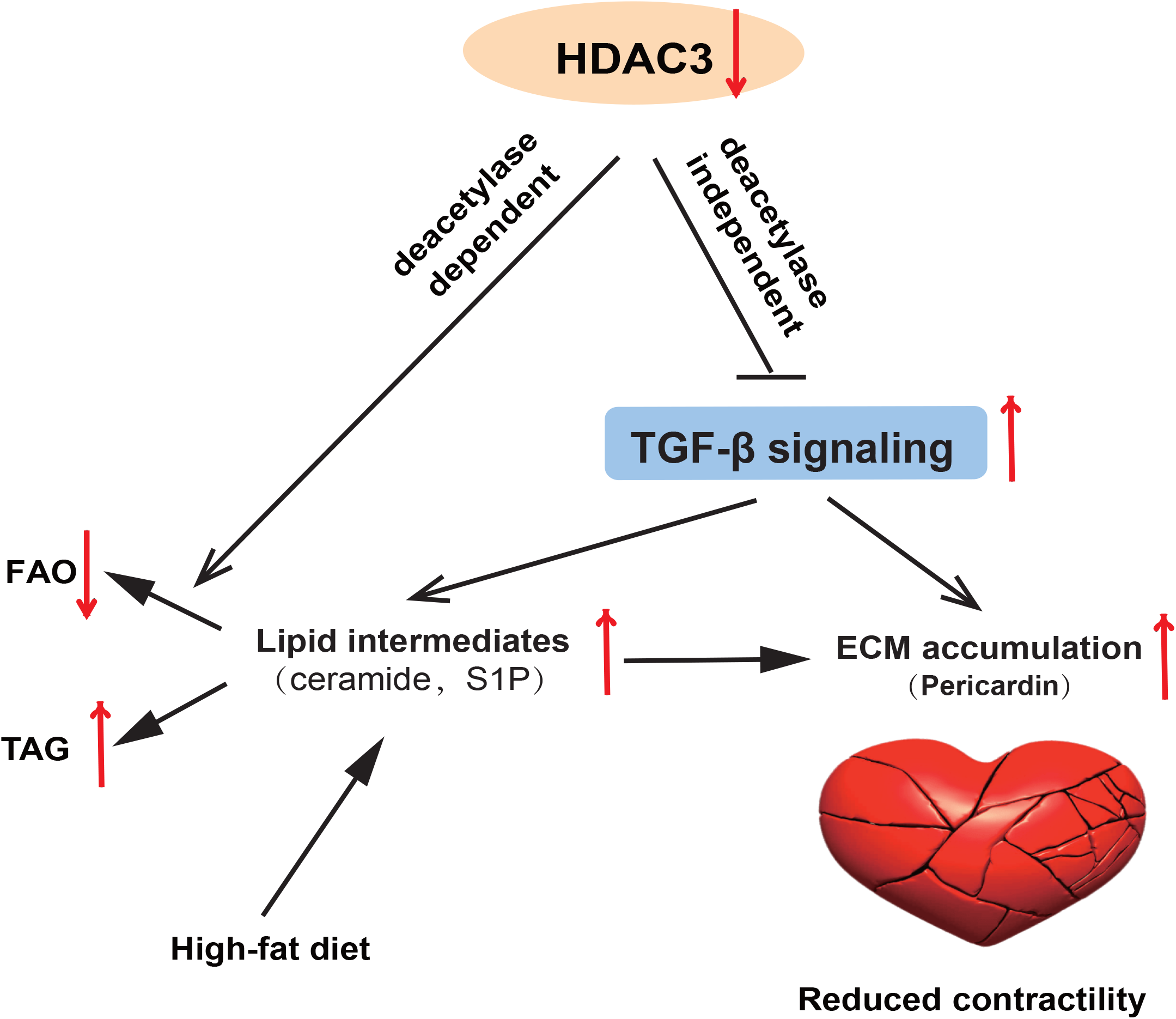
The deacetylase-dependent and deacetylase-independent role in maintaining cardiac performance. HDAC3 play deacetylase-dependent role in lipid homostasis by promoting fatty acid oxidation. The defects of fatty acid oxidation in HDAC3 KD would reroute the lipid intermediates to TAG synthesis. The increase of lipid intermediates S1P would promote ECM accumulation. The non-enzymatic function of HDAC3 would silence the TGF-β signaling to directly prevent ECM accumulation. Additionally TGF-β signaling would indirectly promote cardiac fibrosis through increase the lipid intermediates SIP.

The lipotoxicity promotes the development of cardiomyopathy [30]. Deacetylase dead mutant K26A failed to recover TAG level but succeeded in improving cardiac function (Fig. 1E, Fig. 2D, Fig. 3), indicating TAG accumulation is not equivalent to lipotoxity. Several studies found that the lipid intermediates instead of TAG caused lipotoxity [28, 31, 32]. Knockout of HSL with more TAG accumulation failed to show toxicity to cardiac function [32]. The elevated TAG levels in HDAC3 KO mice is not enough to cause lipotoxicity in liver [28]. In addition, the lipid intermediate sphingosine-1-phosphate (S1P) has been demonstrated to promote the fibrotic development in heart [33]. TGF-β signaling would promote fibrosis through stimulating SphK1 activity that catalyze ceramide to generate S1P [33]. So we speculated that HDAC3 prevent cardiac fibrosis through inhibiting TGF-β signaling and reducing the lipid intermediates by increasing fatty acid oxidation (Figure. 8). It will be interesting to compare the lipid composition between the K26A rescue mutant and HDAC3 knock down to find the true biomarker for HDAC3 cardiac lipotoxicity.

High fat diet is a high risk factor in causing cardiac dysfunction [25]. HDAC3 KD mice displayed severe cardiomyopathy upon switching to a HFD [6]. Our result showed that hearts with HDAC3 KD did not exhibit a more serious cardiac contractility defect nor more increased Pericardin expression upon HFD feeding, even though TAG levels were further increased by HFD in HDAC KD flies.Our results suggest that HDAC3 probably mediates the HFD-induced effects on cardiac function. Cardiac HDAC3 KD in the presence of lipid overload caused severe cardiac contractile dysfunction [6]. Furthermore, a recent study showed that NADPH inhibits HDAC3-DAD domain complex formation by competing with IP4 [34]. Thus, metabolic state may participate in maintain cardiac performance through mediating HDAC3 activity.

## Material & Method

### Fly Stocks

HDAC3 RNAi line (Short Hairpin ID: SH00372.N) and UAS-control line (BL35787) were obtained from the Bloomington Drosophila Stock Center. A second HDAC3 RNAi line (Short Hairpin ID: SH03161.N) was obtained from the stock center of Tsinghua University. The heart specific driver hand 4.2 Gal4, described previously [35], was used to drive HDAC3 short hairpin RNA interference fragment expression. Flies were reared and maintained on a standard cornmeal– yeast diet at 25°C. For high fat diet experiments, we collected 0-5 day old newly enclosed flies that were split into two groups, one fed the standard cornmeal-yeast diet, normal food (NF), and one fed a high fat diet (HFD) consisting of the NF plus 30% coconut oil. Collected flies were maintained on NF or HFD for 5 days at 21°C.

### Intravital imaging of *Drosophila* heart

A transgenic heart marker, R94C02::tdTomato was utilized for heart beat movie[17]. Adult flies were anesthetized using FlyNap (Carolina) and then glued to a coverslip through their dorsal side using optical adhesive glue (Noland #61). Heart beating of individual flies was recorded through the dorsal cuticle with a digital camera (Hamamatsu, ORCA-flash4.0LT, C11440) at 280 frames/seconds. Data were captured using HC Image software (Hamamatsu)

### Semi-intact Optical Heart Function Analysis

*Drosophila* semi-intact heart preparations were prepared as described previously [36]. Movies of beating hearts were recorded for 30 seconds with a high-speed EM-CCD camera (Hamamatsu, C9300) at 130 frames/seconds. Data were captured using HC Image software (Hamamatsu). Movies were analyzed with Semi-automatic Optical Heartbeat Analysis software to quantify heart periods, systolic and diastolic intervals, systolic and diastolic diameters, fractional shortening, and arrhythmia indexes (defined as the standard deviation of the heart period normalized to the median of each fly) and to produce M-mode records [18, 37].

### Immunohistochemical staining of Drosophila heart

Semi-intact Drosophila hearts were prepared and fixed as described previously [38]. Hearts were stained with mouse monoclonal antibody against Pericardin (Developmental Studies Hybridoma Bank (DSHB), #EC11) followed by Cy3 tagged secondary antibody (Jackson ImmunoResearch #144931) or fluorescently-tagged phalloidin (Cell Signaling, # 8878S 330mM final concentration in PBS) directly. Stained hearts were imaged by confocal microscope. Pericardin fluorescent signal of the distal heart is quantified using image J software.

### Triglyceride Assay

Female flies were starved on filter paper soaked in distilled water for 30 min and then placed into Eppendorf tubes for immediate quantification or frozen at -80°C for later processing. 3 female flies per sample were homogenized in 300ul ethanol and then spun at 4000 g on centrifuge for 15 min. 200ul supernatant was transferred into new EP tube and used for conducting TAG assay (Nanjing jiancheng in China, #A110-1-1). The leftover were kept and added 200 ul PBS. Then the mixture was spun at 4000 g on centrifuge for 15 min. Supernatant was transferred into new EP tube and used for conducting Bradford protein assay (Solarbio, #PC0020). Relative TAG content was normalized to protein level.

For cardiac TAG assay, 20 female hearts per sample was dissected in artificial hemolymph and put into 100ul ethanol. After homogenization and centrifuge, 70ul supernatant was transferred into new EP tube and used for conducting TAG assay. The leftover were kept and added 60 ul PBS for conducting Bradford protein assay.

### Western Blots

(A) For each lane 10 females hearts were harvested and directly lysed in 10ul RIPA buffer. Protein was separated by 10% SDS-PAGE gel and transferred to a PVDF membrane. The membrane was then probed with anti-pericardin(DSHB, #EC11) and anti-actin (Invitrogen, #MA5-11869) antibodies, followed by horseradish peroxidase conjugated secondary antibodies (Absin, #20001). Protein bands were visualized using enhanced chemiluminescence reagent (Beyotime, #P0018AS). Pericardin band intensity relative to actin was quantified with Image J software.

### Real time PCR

Total RNA was extracted from 15 female hearts of each genotype with Trizol reagent (Ambion #15596018). The samples were treated with DNase I (Qiagen) to remove DNA. Reverse transcription was performed using the reverse transcription kit (Thermo Scientific, #k1622) according to the manufacturer’s instruction. SYBR Green (Bimake, #B21202) was used as fluorescent dye for qPCR reaction. Thermal cycling and fluorescence monitoring were performed in StepOne Plus Real Time PCR instrument (Applied Biosystems). Primers used are listed below.

Primers:

HDAC3-F: 5’TGAACTACGGACTGCACAAGAA3’

HDAC3-R: 5’CTTCGTATAGGCCACGGAATTG3’

### Survivorship

The progeny of hand Gal4 x UAS-control and hand Gal4 x UAS-HDAC3 RNAi crosses were collected and divided into two populations: one fed with NF and one fed with HFD. Flies were maintained at 21°C, and housed separately in groups of 25 flies per vial. Around 180 flies were prepared, and flipped to fresh food every 3^rd^ day; survival was scored following each transfer. Lifespan data is presented as Kaplan-Meier plots, and significance was determined using Mandel-Cox Log rank test by Graphpad Software.

### Statistics

All statistical analysis was performed using GraphPad Prism version 8.0 (GraphPad Software, San Diego California USA); Specific tests used are indicated in the figure legends.

## Acknowledgments

We thank Mattias Mannervik for reagents and useful discussions.

## Supplemental materials

**Supplemental Figure 1. HDAC3 RNA from cardiac HDAC3 KD hearts in qPCR**

Relative expression of HDAC3 was measured in 1-week-old adult female hearts (15 hearts/sample) and was normalized to rp49 expression. Cardiac-specific HDAC3 KD showed 47.7% reduction compared to control. Significance was determined using an unpaired Student’s T-test. Differences relative to the control are indicated by individual asterisks; GraphPad statistical analysis, **p<0.01.

**Supplemental Figure 2. Cardiac-specific HDAC3 KD using a second HDAC3 RNAi line also causes cardiac dysfunction**.

(A) Heart Period, (B) Systolic Interval, (C) Diastolic Interval, (D) contractility quantified as Fractional Shortening, (E) Systolic Diameter, and (F) Diastolic Diameter were measured for hearts from wild type control flies (Hand/+), and flies with cardiac specific HDAC3 knockdown (Hand>UAS-HDAC3 RNAi, SH03161.N). Note the significant systolic interval prolongation and decreased contractility in HDAC3 KD. Significance was determined using unpaired student’s t tests. Differences relative to the control are indicated by individual asterisks; Graph Pad statistical analysis, **P<0.01, ns represents no significance. Sample size was 20 to 30 flies per genotype.

## Supplemental movies

**Movie S1**. Movie of a Semi-Intact Heart Preparation from control flies.

**Movie S2**. Movie of a Semi-Intact Heart Preparation from HDAC3 KD flies.

