## Supplementary figures and images for "The deacetylase dependent and independent role of HDAC3 in cardiomyopathy"

### Fig. S1

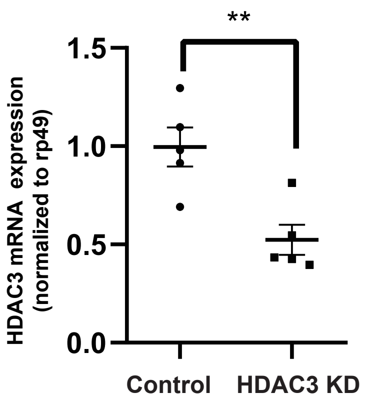

### Fig. S2

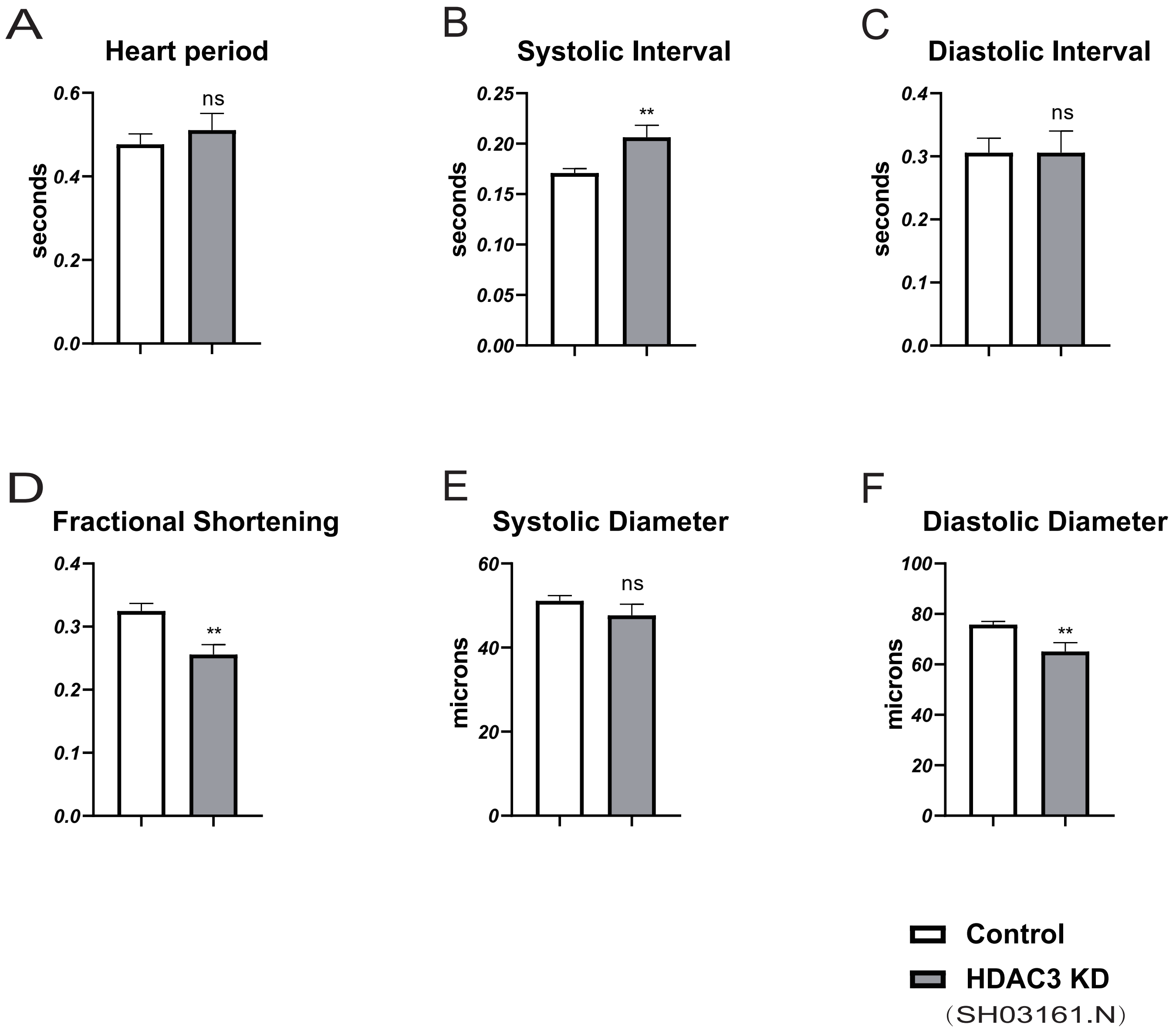
